# Nitrate inhibits nodule organogenesis through inhibition of cytokinin biosynthesis in *Lotus japonicus*

**DOI:** 10.1101/2020.11.03.366971

**Authors:** Jieshun Lin, Yuda Purwana Roswanjaya, Wouter Kohlen, Jens Stougaard, Dugald Reid

## Abstract

Legumes balance nitrogen acquisition from soil nitrate with symbiotic nitrogen fixation. Nitrogen fixation requires establishment of a new organ, which is a cytokinin dependent developmental process in the root. We found cytokinin biosynthesis is a central integrator, balancing nitrate signalling with symbiotic acquired nitrogen. Low nitrate conditions provide a permissive state for induction of cytokinin by symbiotic signalling and thus nodule development. In contrast, high nitrate is inhibitory to cytokinin accumulation and nodule establishment in the root zone susceptible to nodule formation. This reduction of symbiotic cytokinin accumulation was further exacerbated in cytokinin biosynthesis mutants, which display hypersensitivity to nitrate inhibition of nodule development, maturation and nitrogen fixation. Consistent with this, cytokinin application can rescue nodulation and nitrogen fixation of biosynthesis mutants in a concentration dependent manner. These inhibitory impacts of nitrate on symbiosis occur in a *Nlp1* and *Nlp4* dependent manner and contrast with the positive influence of nitrate on cytokinin biosynthesis that occurs in non-symbiotic species. Altogether this shows that legumes, as exemplified by *Lotus japonicus*, have evolved a different cytokinin response to nitrate compared to non-legumes.

**One sentence summary:** Cytokinin biosynthesis is suppressed by nitrate in *Lotus japonicus*, providing a mechanism for nitrate inhibition of symbiotic nodule organogenesis.

## Introduction

Nitrogen deficiency is the most common nutritional limitation to plant growth. Legumes can overcome this limitation by acquiring nitrogen through the establishment of a symbiotic relationship with nitrogen fixing rhizobia. The establishment of a new organ (a nodule) and transfer of resources to the bacterial partner makes symbiotic nitrogen fixation less favourable than uptake of soil nitrate. Therefore, nitrate is preferentially acquired and nodule development is inhibited in soils with high nitrate levels (Oldroyd and Leyser, 2020).

Nodule development is initiated through a common symbiotic pathway shared with establishment of arbuscular mycorrhizal fungus (Kistner and Parniske, 2002; Madsen et al., 2010; Martin et al., 2017). A major downstream target in the establishment of nodules is the induction of cytokinin synthesis and signalling (Gonzalez-Rizzo et al., 2006; Murray et al., 2007; Tirichine et al., 2007; Reid et al., 2017; van Zeijl et al., 2015). Cytokinin is essential for nodule organogenesis, as exemplified by loss of nodule development in cytokinin receptor mutants (Gonzalez-Rizzo et al., 2006; Murray et al., 2007; Held et al., 2014; Boivin et al., 2016). Stimulation of cytokinin signalling through biosynthesis and receptor activation is also sufficient to trigger nodule development, even in the absence of rhizobia (Tirichine et al., 2007; Heckmann et al., 2011; Reid et al., 2017; Liu et al., 2018). This cytokinin signalling regulates expression of central components of nodulation signalling, including the transcription factor NIN (Nodule Inception; Schauser et al., 1999) via distal *cis*-elements in the *Nin* promoter (Liu et al., 2019). Cytokinin- and NIN-dependent signalling also initiates negative feedback of nodule organogenesis and infection via induction of CLE (CLAVATA3/ESR-related) peptides and the AON (Autoregulation of Nodulation) pathway (Soyano et al., 2014; Laffont et al., 2020; Ferguson et al., 2019). The AON pathway integrates signals from both prior nodulation events and soil nitrate availability to balance nodule development with plant resources.

Nitrate signalling depends on uptake and perception by nitrate transceptors (transporter-receptors) of the NRT1.1 family (Ho et al., 2009; Bouguyon et al., 2015). The majority of transcriptional responses to nitrate are then controlled by the action of NIN-Like proteins (NLPs) (Castaings et al., 2009; Liu et al., 2017). In legumes, NLP signalling is tightly integrated with symbiotic signalling with NLPs regulating symbiotic signalling both directly and through competition for NIN binding sites (Lin et al., 2018; Nishida et al., 2018). Loss-of-function mutants in legume NLPs (eg. *Ljnrsym1/Ljnlp4* and *Mtnlp1*) therefore show nitrate resistant symbiosis phenotypes.

In addition to local responses to nitrate, plants possess systemic regulatory circuits allowing response to nitrate to be coordinated between roots and shoots. Cytokinin signalling is one of the pathways underlying this systemic signalling of nitrate availability (Sakakibara, 2020). Outside of legumes, induction of cytokinin biosynthesis by nitrate has been described in several species including maize (Takei et al., 2001), rice (Kamada-Nobusada et al., 2013) and Arabidopsis (Miyawaki et al., 2004). Several cytokinin biosynthesis (IPT, LOG and CYP735A enzymes) and transport (ABCG14) components have subsequently been shown to coordinate plant responses to nitrogen. For example, *AtIPT3* and *AtABCG14* activity in the root vasculature can increase cytokinin export in response to nitrate supply (Miyawaki et al., 2004; Zhang et al., 2014; Poitout et al., 2018), while *AtIPT3* and *AtCYP735A2* have been implicated in NLP dependent nitrate signalling (Maeda et al., 2018; Takei et al., 2004a, 2004b). In rice, four IPT genes and a resulting accumulation of cytokinin were identified as nitrate and ammonium responsive (Kamada-Nobusada et al., 2013).

Given the positive role of cytokinin in nodule development, in contrast to the inhibitory role of nitrate in this process, it remains to be seen how and whether cytokinin synthesis and signalling play equivalent signalling and coordination roles in nitrogen signalling in legumes. Here, we investigate this link and find nitrate is inhibitory to cytokinin biosynthesis in the model legume *Lotus japonicus.* This provides a regulatory target for nitrate signalling, ensuring nodule development and soil nitrate is balanced and implies additional regulatory mechanisms must be recruited or amplified in legumes to signal nitrogen availability.

## Results

### Rhizobia induced cytokinin biosynthesis is inhibited by nitrate

In response to rhizobia, *L. japonicus* induces the expression of cytokinin biosynthesis genes, including *Ipt2; Log1;* and *Log4*, to trigger nodule organogenesis (Reid et al., 2017). This induction occurs primarily in the region of emerging root hairs at the root tip, known as the susceptible zone (Figure 1a). While environmental nitrate induces cytokinin synthesis in many species, in legumes high nitrate can inhibit nodule organogenesis, raising the question of how legumes deal with this apparent paradox? To investigate the effect of nitrate on cytokinin biosynthesis during nodule initiation, we first analysed the expression of cytokinin biosynthesis genes. In the absence of nitrate, *Ipt2; Ipt3; Ipt4; Log1;* and *Log4* are up-regulated in the susceptible zone following rhizobia inoculation, in line with previous results (Fig 1A-D; S1) (Reid et al., 2017). However, in the presence of high nitrate which is inhibitory to nodule initiation (5 mM KNO_3_), the relative transcript abundance of *Ipt2; Ipt3; Ipt4; Log1;* and *Log4* was lower at both one and two days post inoculation (dpi), relative to plants grown in the absence of nitrate (Fig 1B-D; S1).

**Figure 1.**
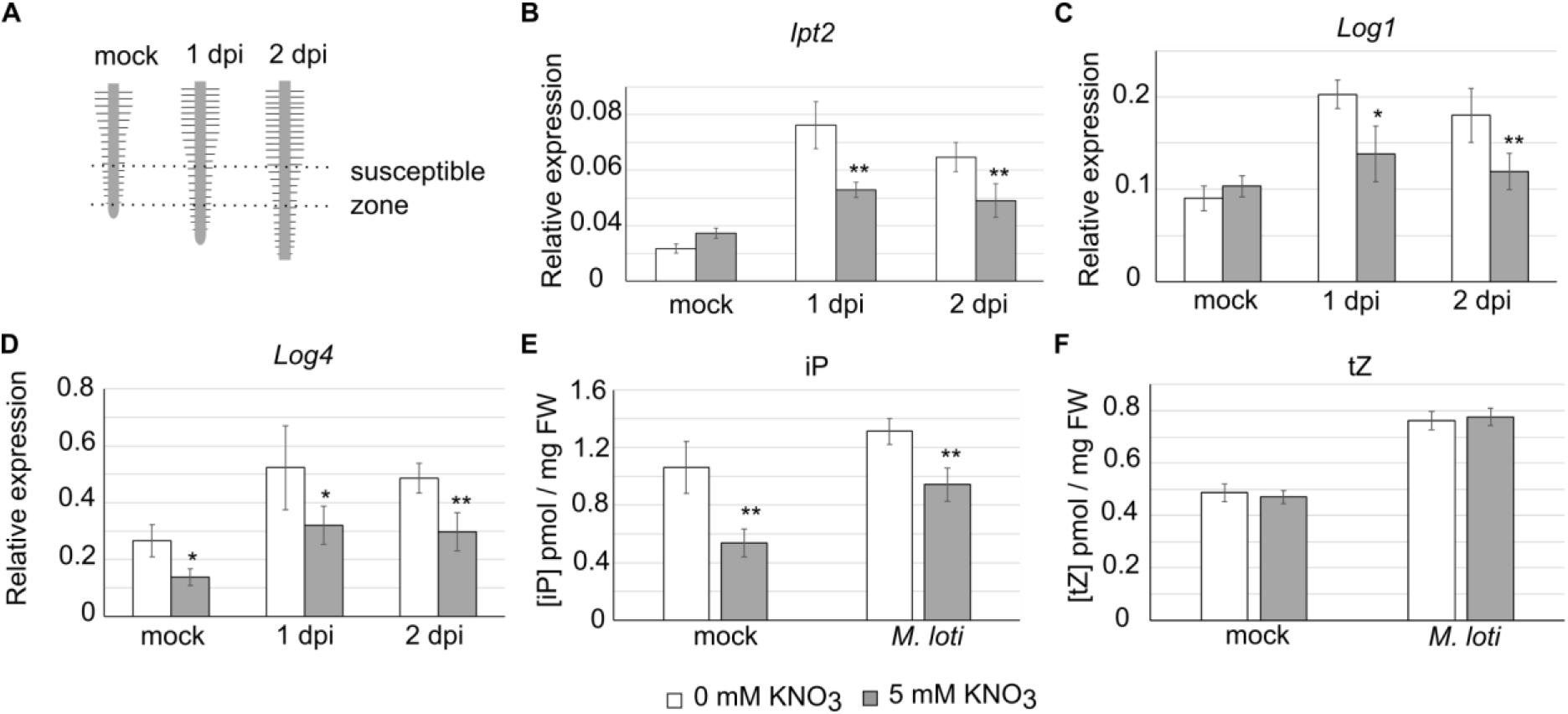
Nitrate inhibits induction of symbiotic cytokinin biosynthesis. A, The root zone susceptible to nodule initiation used for qRT-PCR and cytokinin analysis is indicated. B-D, Relative transcript abundance by qRT-PCR 1 and 2 days post inoculation (dpi) with *M. loti* in absence and presence of 5 mM KNO_3_. E-F, Cytokinin free base content (iP and *t*Z) analysed in mock or 2 days post inoculation with*M. loti* in absence and presence of 5 mM KNO_3_. Ubiquitin is used as a reference gene. Bars show mean +/-SE for *n*=5 in qRT-PCR and *n*=6 in cytokinin analysis. Significant differences between nitrate presence and absence conditions is indicated by *<0.05 and **<0.01 as determined by Student’s t-test.

To confirm that this nitrate-induced suppression of transcript levels resulted in altered levels of cytokinin, we quantified the cytokinin ribosides and bases in the same tissue and conditions two days post inoculation. We found that both iP and *t*Z cytokinin bases are induced by rhizobia inoculation in both the absence and presence of nitrate (Fig 1E,F). In agreement with the gene expression analysis, rhizobia inoculation results in an increase in iP levels, while nitrate exposure significantly reduces iP relative to nitrate-free plants irrespective of inoculation status (Fig 1E). The levels of *t*Z were not significantly different in the two nitrate conditions (Fig 1F).

### Cytokinin biosynthesis mutants show reduced iP content in N sufficient conditions

Given that iP production is inhibited by nitrate, we hypothesised that *ipt* biosynthesis mutants may exacerbate this reduction. We therefore analysed the cytokinin content of *ipt3-1, ipt4-1* and the *ipt3-2 ipt4-1* double mutant two days post rhizobia inoculation in the absence and presence of nitrate (Fig 2). In uninoculated conditions, all of these mutants showed reduced iP content in the susceptible zone relative to the wild-type Gifu. Similar to the wild-type, in all cases iP content increased after inoculation, likely through activity of the symbiotic responsive *LjIpt2* (Reid et al., 2017) and confirming the redundancy present in the cytokinin biosynthesis pathway. In high nitrate both the *ipt4-1* and *ipt3-2 ipt4-1* mutants showed reduced iP content after inoculation relative to plants grown without nitrate. Under high nitrate, the *ipt4-1* and *ipt3-2 ipt4-1* mutants showed the lowest iP content following inoculation with 54.9% and 46.3% of wild-type levels in the same condition.

**Figure 2.**
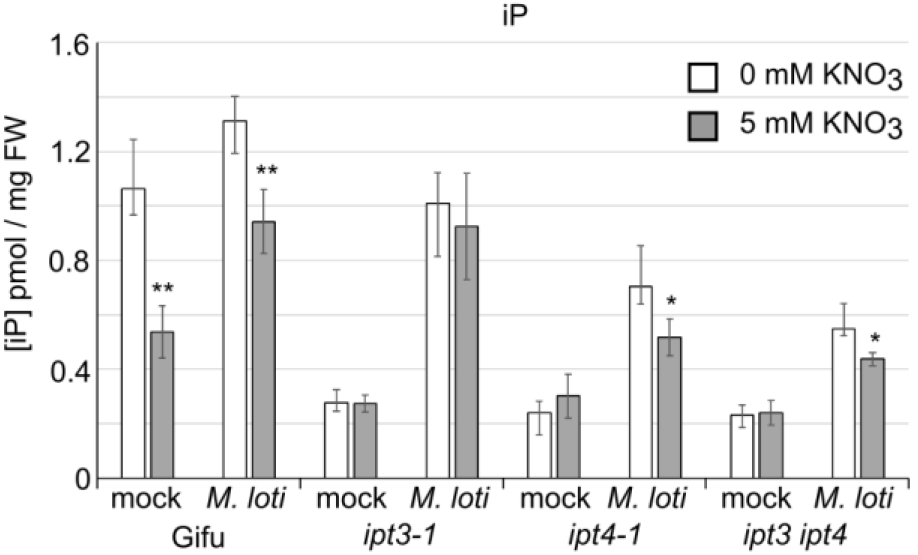
*ipt3* and *ipt4* mutants show reduced iP content in high nitrate conditions. Cytokinin iP free base content analysed in mock or 2 days post inoculation with *M. loti* in absence and presence of 5 mM KNO_3_ is shown. Bars show mean +/− SE for n=6. Significant differences between nitrate presence and absence conditions is indicated by *<0.05 and **<0.01 as determined by Student’s t-test.

### Cytokinin biosynthesis mutants are hyper-sensitive to nitrate inhibition of nodule organogenesis but not infection

To assess the impact of the reduced iP on nodule development, we investigated the nodulation phenotypes of *ipt3-1, ipt4-1* and *ipt3-2 ipt4-1* relative to Gifu in the absence or presence of 5 mM nitrate. In the absence of nitrate, *ipt3-1* and *ipt4-1* mutants do not show significantly reduced nodulation while *ipt3-2 ipt4-1* formed slightly fewer nodules in these conditions (Fig 3A-C). In high nitrate, nodule initiation (indicated here as total nodules) and nodule maturation (as determined by nodules acquiring a distinct pink-red colour) are both significantly impaired on *ipt3-1, ipt4-1* and *ipt3-2 ipt4-1* roots (Fig 3A-C). In particular, on *ipt3-2 ipt4-1* roots, only small bumps that do not fully mature are observed (Fig 3A).

**Figure 3.**
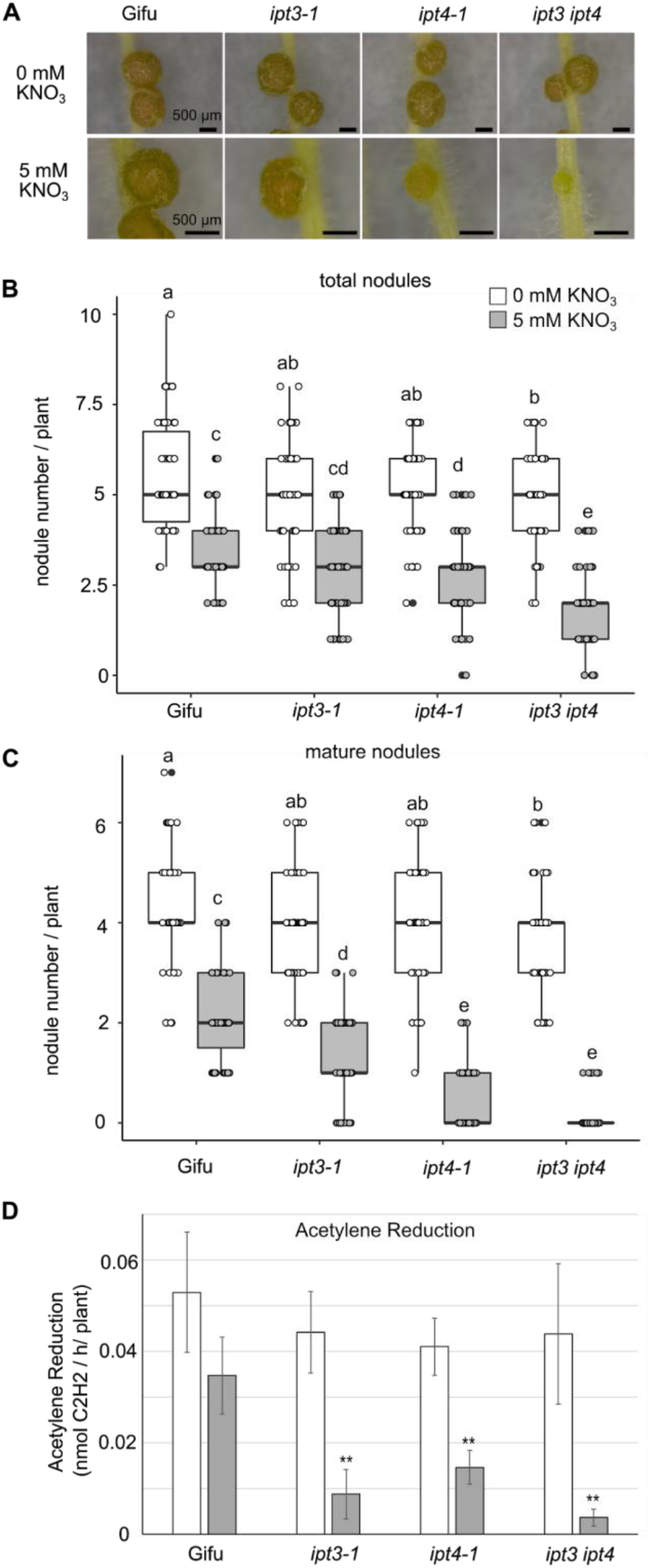
*Ipt3* and *Ipt4* are required for resistance to nitrate inhibition of nodulation. A, Images of nodules developed in the absence and presence of 5 mM KNO_3_ at 14 dpi with *M. loti* on the indicated host genotypes. Scale bar = 500 μM. B-C, Development of pink (B) and total (C) nodules in the absence and presence of 5 mM KNO_3_ at 14 dpi with *M. loti.* D, Nitrogenase activity assessed by Acetylene Reduction Assay (ARA) at 21 dpi with *M. loti* in the absence and presence of 5 mM KNO_3_. n≥50 in nodulation assay, significant differences among different genotypes and nutrient conditions are indicated by letters (p<0.05) as determined by ANOVA and Tukey post-hoc testing. n=6 in ARA, significant differences between Gifu and mutants are indicated by **<0.01 as determined by Student’s t-test.

The impact of impaired nodule maturation on nitrogen fixation was assessed by Acetylene Reduction Assay (ARA). In line with our scoring of nodule development, in the absence of nitrate, there is no significant difference in ARA in these mutants. Under high nitrate conditions, Gifu shows a significant 35.4% reduction in ARA activity relative to nitrate free conditions (Fig 3D). We found *ipt3-1, ipt4-1* and *ipt3-2 ipt4-1* mutants all showed a significantly greater sensitivity to nitrate inhibition, exhibiting 80.2%, 64.3% and 91.7% reduction in ARA respectively relative to nitrate free conditions (Fig 3D).

To study whether *Ipt3* and *Ipt4* also play a role in resistance to nitrate inhibition of early nodule initiation and rhizobia infection (IT), we counted the nodule primordia and infection threads (IT) that developed 7 dpi with *M. loti.* In the absence of nitrate, *ipt3-2 ipt4-1* does not form significantly less nodule primordia than Gifu. However, in high nitrate conditions, *ipt3-2 ipt4-1* did not develop any nodule primordia at 7dpi, while a few visible primordia are already developed on Gifu (Fig 4A).

**Figure 4.**
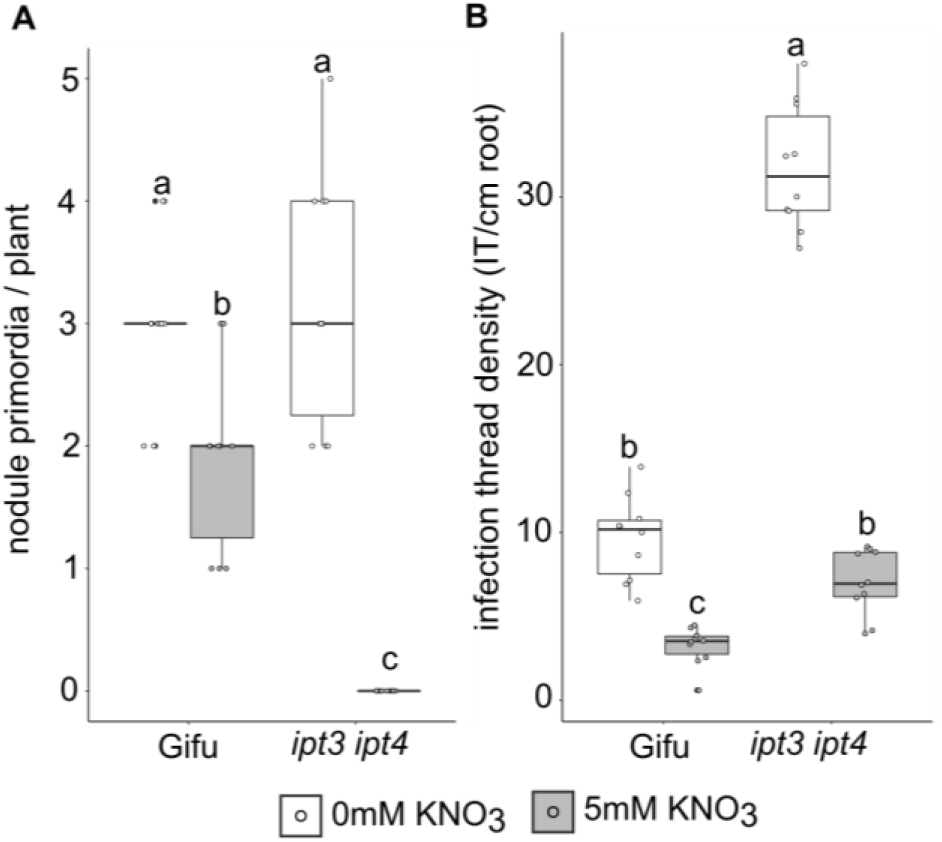
*Ipt3* and *Ipt4* contribute to nitrate resistance to nodule initiation but not infection thread formation. A, Nodule primordia number in absence and presence of 5 mM KNO_3_ at 7dpi with *M. loti.* B, Infection thread density in absence and presence of 5 mM KNO_3_ at 7dpi with *M. loti.* n=10, significant differences between genotypes and nutrient conditions are indicated by letters (p<0.05) as determined by ANOVA and Tukey post-hoc analysis.

In contrast to the positive role in nodule organogenesis, cytokinin plays a negative role in regulating infection by rhizobia (Murray et al., 2007). Similar to the hyperinfection phenotype of the *lhk1* cytokinin receptor mutants (Murray et al., 2007), *ipt3-2 ipt4-1* forms more than 3 times more IT than Gifu, implying cytokinin biosynthesis plays a negative role in IT formation (Fig 4B). However, under high nitrate conditions, both Gifu and *ipt3-2 ipt4-1* show significantly reduced IT formation relative to nitrate free plants (Fig 4B).

### Cytokinin application rescues nitrate inhibition of nodule initiation and maturation

Because IPTs are key enzymes in cytokinin biosynthesis, we asked whether application of the cytokinin 6-Benzylaminopurine (BA) could rescue the hypersensitivity to nitrate inhibition seen in *ipt3-2 ipt4-1.* When grown in the presence of 10^−8^ M BA, Gifu develops shorter roots and forms less nodules compared with mock treatment (Fig 5A-C and Fig S2). This inhibition of root elongation is also seen in *ipt3-2 ipt4-1* grown on 10^−8^ M BA (Fig S2). However, grown in the presence of BA, *ipt3-2 ipt4-1* can form on average 0.47 (10^−9^ M BA) and 1.15 (10^−8^ M BA) mature pink nodules under high nitrate condition, which is otherwise completely inhibitory to mature nodule formation (Fig 5A-C). Applying BA at lower concentration (10^−9^ M BA) is also able to trigger formation of more total nodules on *ipt3-2 ipt4-1* in the presence of nitrate, while 10^−8^ M BA did not increase total nodule numbers (Fig 5B).

**Figure 5.**
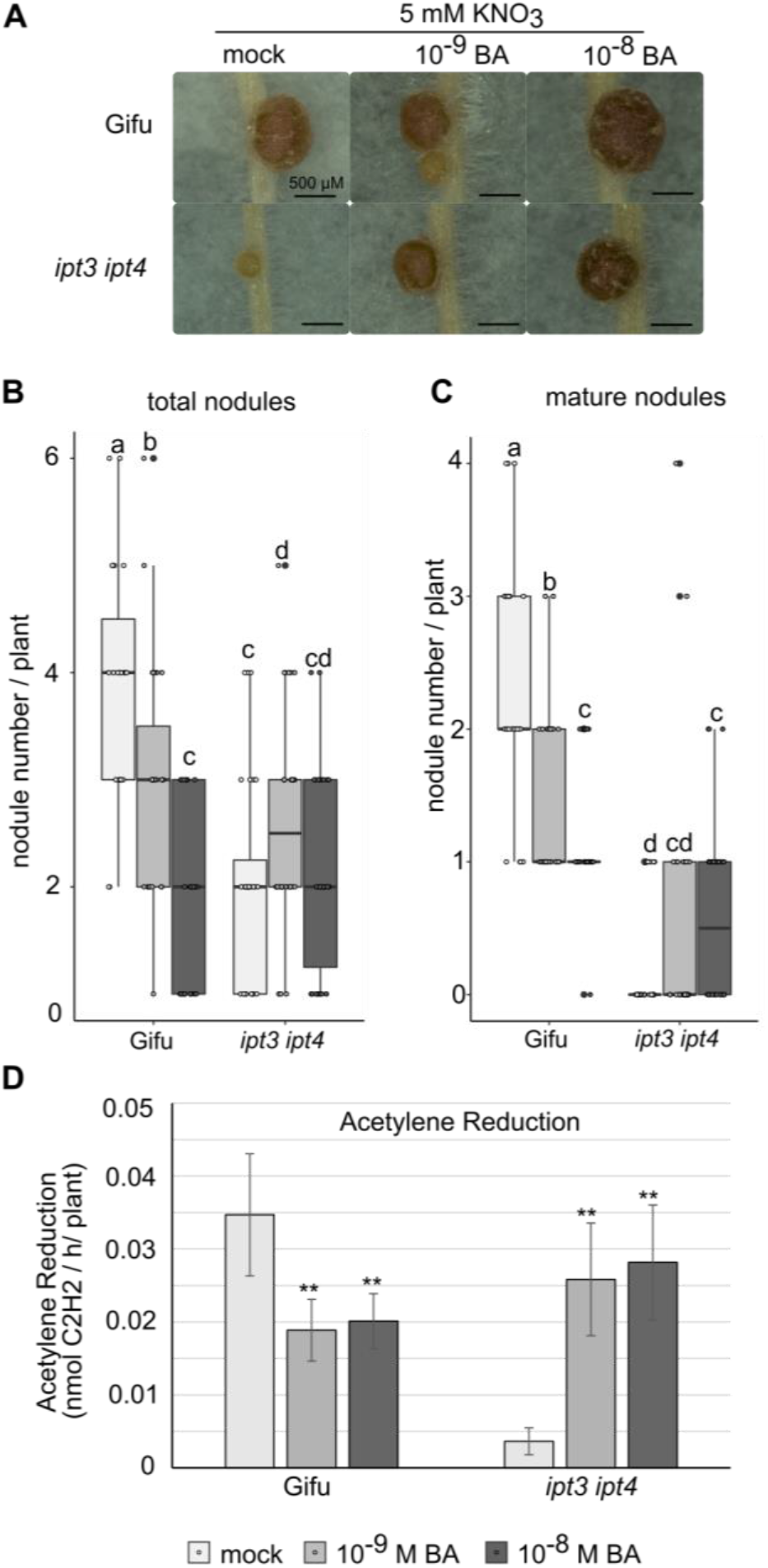
Nitrate inhibition of nodule development can be rescued by cytokinin application.A, Images of nodules developed in the presence of 5 mM KNO_3_ with mock, 10^−9^ or 10^−8^ M BA at 14 dpi with *M. loti* on the indicated host genotypes. Scale bar = 500 μM. B-C, Development of pink (B) and total (C) nodules in the presence of 5 mM KNO_3_ with mock, 10^−9^ or 10^−8^ M BA at 14 dpi with *M. loti.* D, Nitrogenase activity assessed by ARA at 21 dpi with *M. loti* in the presence of 5 mM KNO_3_ with mock, 10^−9^ or 10^−8^ M BA. n≥25 in nodulation assay, significant differences among different genotypes and concentration of BA are indicated by letters (p<0.05) as determined by ANOVA and Tukey post-hoc testing. n=6 in ARA, significant differences between mock and BA application are indicated by **<0.01 as determined by Student’s t-test.

We also measured nitrogenase activity by ARA in the BA rescued plants. Consistent with the pink nodule number, the ARA activity of BA treated *ipt3-2 ipt4-1* is significantly higher than untreated plants in high nitrate conditions (Fig 5D). Cytokinin application is thus able to rescue nitrate inhibition of nodule initiation, maturation and nitrogen fixation of the *ipt3-2 ipt4-1* mutant.

### Nitrate inhibition of cytokinin biosynthesis requires *Nlp1* and *Nlp4*

NLP1 and NRSYM1 (here called NLP4) play central roles in nitrate signaling and nitrate inhibition of nodulation (Lin et al., 2018; Nishida et al., 2018), thus we obtained LORE1 insertion lines (Małolepszy et al., 2016) for each gene and characterised their phenotypes (Fig S3A). In the absence of nitrate, *nlp1-2* forms slightly fewer nodules, while *nlp4-1* was not different to wild-type Gifu (Fig S3C-D). In high nitrate, where Gifu forms few pink nodules, *nlp1-2* and *nlp4-1* are able to form significantly more mature pink nodules, which is consistent with previous observations for *nlp* mutants in *M. truncatula* and *L. japonicus* (Lin et al., 2018; Nishida et al., 2018).

To assess the role of *Nlp1* and *Nlp4* mediated signaling in nitrate inhibition of symbiotic cytokinin biosynthesis, we analysed expression of cytokinin biosynthesis genes in the respective mutant backgrounds. As demonstrated in the earlier experiments, *Ipt2, Log1* and *Log4* are all induced after *M. loti* inoculation and suppressed by nitrate (Fig 6). Both *nlp1-2* and *nlp4-1* show induction of *Ipt2, Log1* and *Log4* after *M.loti* inoculation, similar to what is seen in Gifu. However, neither *nlp1-2* nor *nlp4-1* mutants show suppression of these cytokinin biosynthesis genes by nitrate, with *nlp4-1* having slightly higher *Log4* expression (Fig 6).

**Figure 6.**
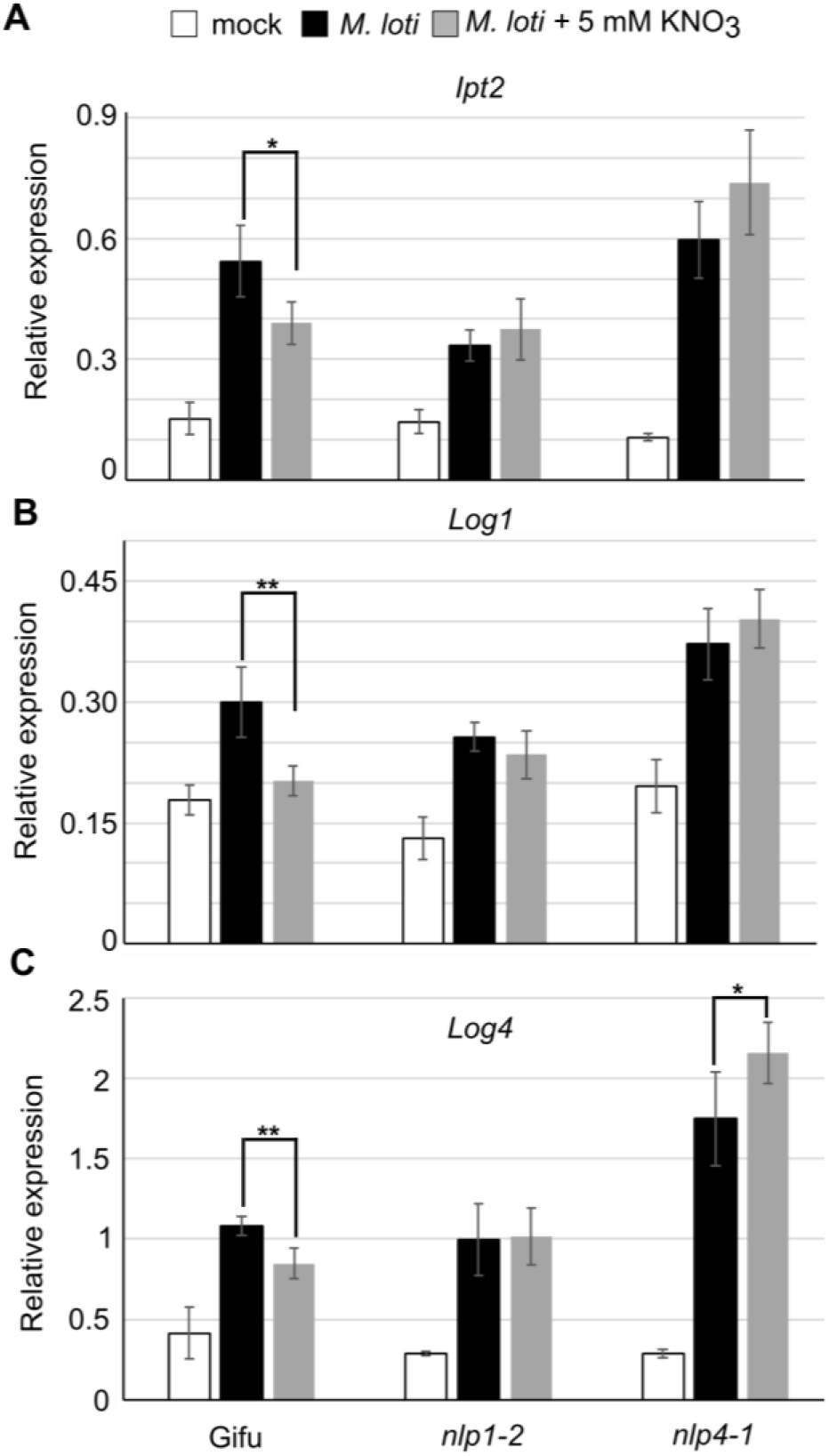
Nitrate inhibition of cytokinin biosynthesis requires *Nlp1* and *Nlp4.* Relative transcript abundance of *Ipt2* (A), *Log1* (B) and *Log4* (C) by qRT-PCR 1 dpi with *M. loti* in absence and presence of 5 mM KNO_3_ in the indicated genotypes. Ubiquitin is used as a reference gene. Bars show mean +/-SE for *n*=5. Significant differences between nitrate presence and absence conditions is indicated by *<0.05 and **<0.01 as determined by Student’s t-test.

Given this insensitivity to nitrate inhibition of cytokinin biosynthesis gene expression, we assessed the ability of *nlp1-2* and *nlp4-1* to rescue the nitrate sensitivity of the *ipt4* mutant. In high nitrate, *ipt4-1* shows significantly impaired nodule maturation with very few mature pink nodules compared with Gifu (Fig 7A-C). However, *ipt4-1 nlp1-2* and *ipt4-1 nlp4-1* double mutants show nitrate resistant nodulation in line with the *nlp1-2* and *nlp4-1* phenotypes, including developing mature nodules (Fig 7A) in increased numbers when compared with *ipt4-1* or Gifu (Fig 7B-C).

**Figure 7.**
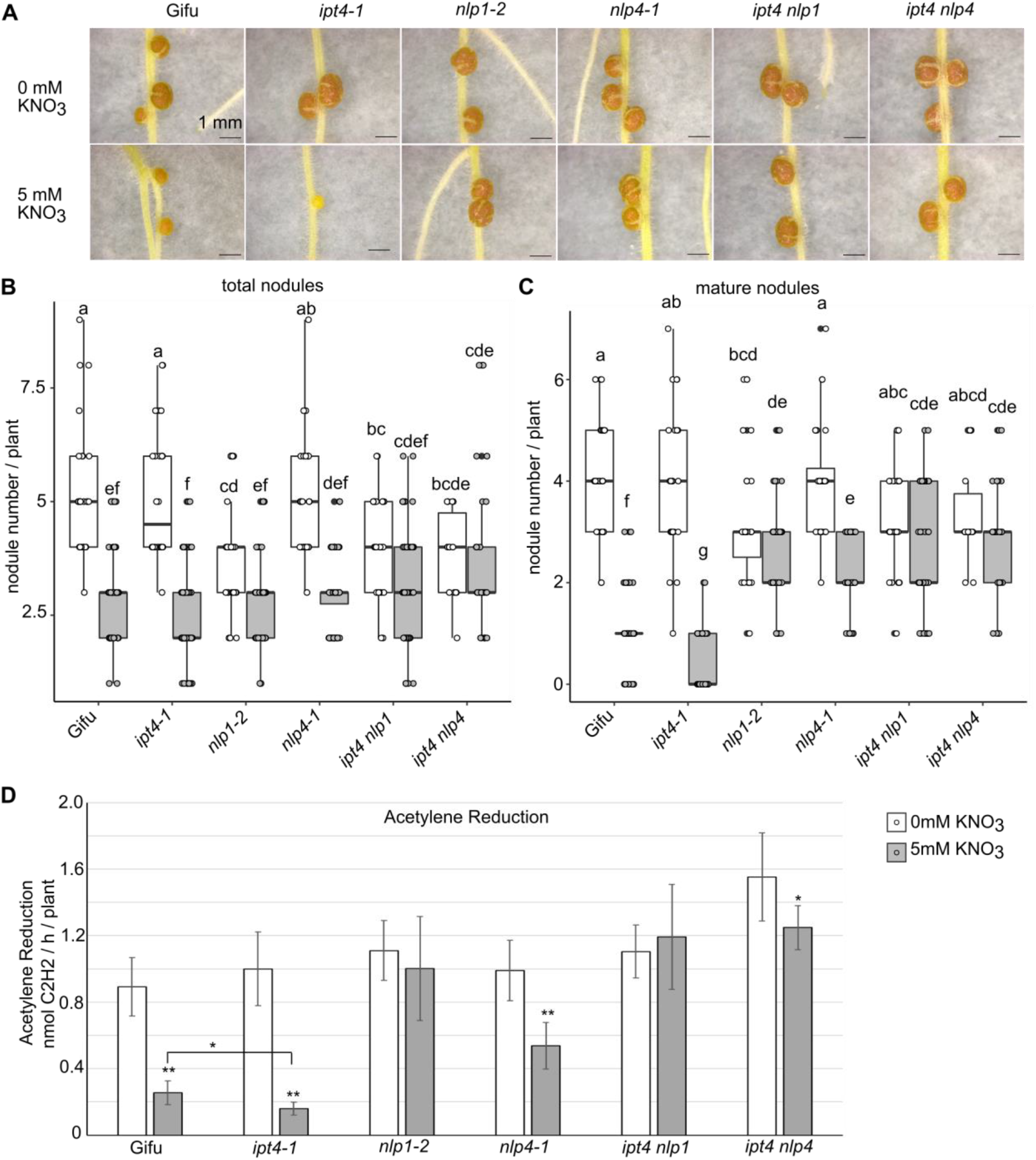
Nitrate signalling mediated by *Nlp1* and *Nlp4* acts upstream of cytokinin biosynthesis to restrict symbiotic signalling. A, Images of nodules developed in the absence and presence of 5 mM KNO_3_ at 14 dpi with *M. loti* on the indicated host genotypes. Scale bar = 1 cm. B-C, Development of pink (B) and total (C) nodules in the absence and presence of 5 mM KNO_3_ at 14 dpi with *M. loti.* D, Nitrogenase activity assessed by ARA at 21 dpi with *M. loti* in the absence and presence of 5 mM KNO_3_. n?25 in nodulation assay, significant differences among different genotypes and nutrient conditions are indicated by letters (p<0.05) as determined by ANOVA and Tukey post-hoc testing. n=6 in ARA, significant differences between Gifu and mutants are indicated by *<0.05 and **<0.01 as determined by Student’s t-test.

To assess the rescue of nodule function by *nlp* mutations, we also measured nitrogenase activity of these mutants by ARA at 21 dpi. In line with the nodule scoring, in the absence of nitrate, there was no significant difference between Gifu and any of the mutants (Fig 7D). In high nitrate conditions, which inhibits nodule ARA activity compared with nitrate-free condition, *ipt4-1* exhibits a further reduction in ARA activity relative to Gifu (Fig 7D). However, *nlp1-2* and *ipt4-1 nlp1-2* maintain equivalent ARA activity to nitrate free conditions. Although *nlp4-1* and *ipt4-1 nlp4-1* exhibit lower ARA activity compared with nitrate-free condition, the nitrate inhibition of ARA activity are significantly less than in Gifu (Fig 7D).

### *Nlp1* and *Nlp4* mediated nitrate signaling inhibit cytokinin biosynthesis via interfering NF signaling

To identify nitrate regulated genes that may contribute to the inhibition of symbiotic cytokinin biosynthesis, we exposed nitrogen starved plants to high levels of nitrate (10 mM KNO_3_) over a time series (0.25 h, 0.5 h, 1 h, 24 h and 72 h) and conducted RNA-seq. Similar to the rapid onset of primary nitrate responses in other species (Krouk et al., 2010), we identified 425 genes that respond to nitrate within 15 minutes, and up to 4411 genes differentially regulated 72 h after nitrate exposure (Fig S4A). Among these genes, many nitrate marker genes, such as *Nrt2.1a, Nrt2.1b, Nia* and *Nir*, are induced at all time points in the series (Fig S4B). On the other hand, nitrogen starvation marker genes such as *Cep1* and *Cep7*, are suppressed after 24 h nitrate exposure (Fig S4B). Taken together, the general nitrate response in *L. japonicus* is similar to other species. However, in contrast to the induction of cytokinin biosynthesis, particularly *AtIPT3* by nitrate in Arabidopsis, none of the *LjIpt* or *LjLog* genes was significantly induced by nitrate across our time series (Fig S5).

Given symbiotic cytokinin biosynthesis is regulated by NF signaling (van Zeijl et al., 2015; Reid et al., 2017), we analysed the expression of key components in early nodulation signalling, including NF signaling and downstream transcription factor genes under nitrate exposure. After 24 h nitrate exposure, *Nfr1, Nfr5, Nsp2* and *Nin* were all suppressed, although the inhibitions of *Nfr1* and *Nfr5* did not persist after 72 h exposure (Fig 8A). To investigate whether *Nlp* mediated nitrate signaling plays a role in the nitrate suppression of these components, we investigated *Nfr1, Nfr5, Nsp2* and *Ern1* expression following nitrate exposure and rhizobia inoculation in Gifu, *nlp1-2* and *nlp4-1.* Consistent with previous RNAseq studies (Larrainzar et al., 2015; Kelly et al., 2018), rhizobia inoculation upregulates *Nsp2* and *Ern1* expression, while *Nfr1* and *Nfr5* expression is suppressed (Fig 8B-E). In turn, expression of *Nfr1, Nfr5, Nsp2* and *Ern1* were all significantly reduced in high nitrate in Gifu (Fig 8B-E). In contrast, *nlp1-2* mutants showed no significant suppression of these genes in the high nitrate condition. In *nlp4-1* mutants, *Nfr1* expression was suppressed by nitrate, while either no reduction *(Nfr5, Ern1)* or even greater induction *(Nsp2)* was found for the other genes in high nitrate conditions (Fig 8B-E). Together, this indicates *Nlp1* and *Nlp4* mediate nitrate signaling repression of the upstream NF perception and signaling components in addition to their suppression of cytokinin biosynthesis expression.

**Figure 8.**
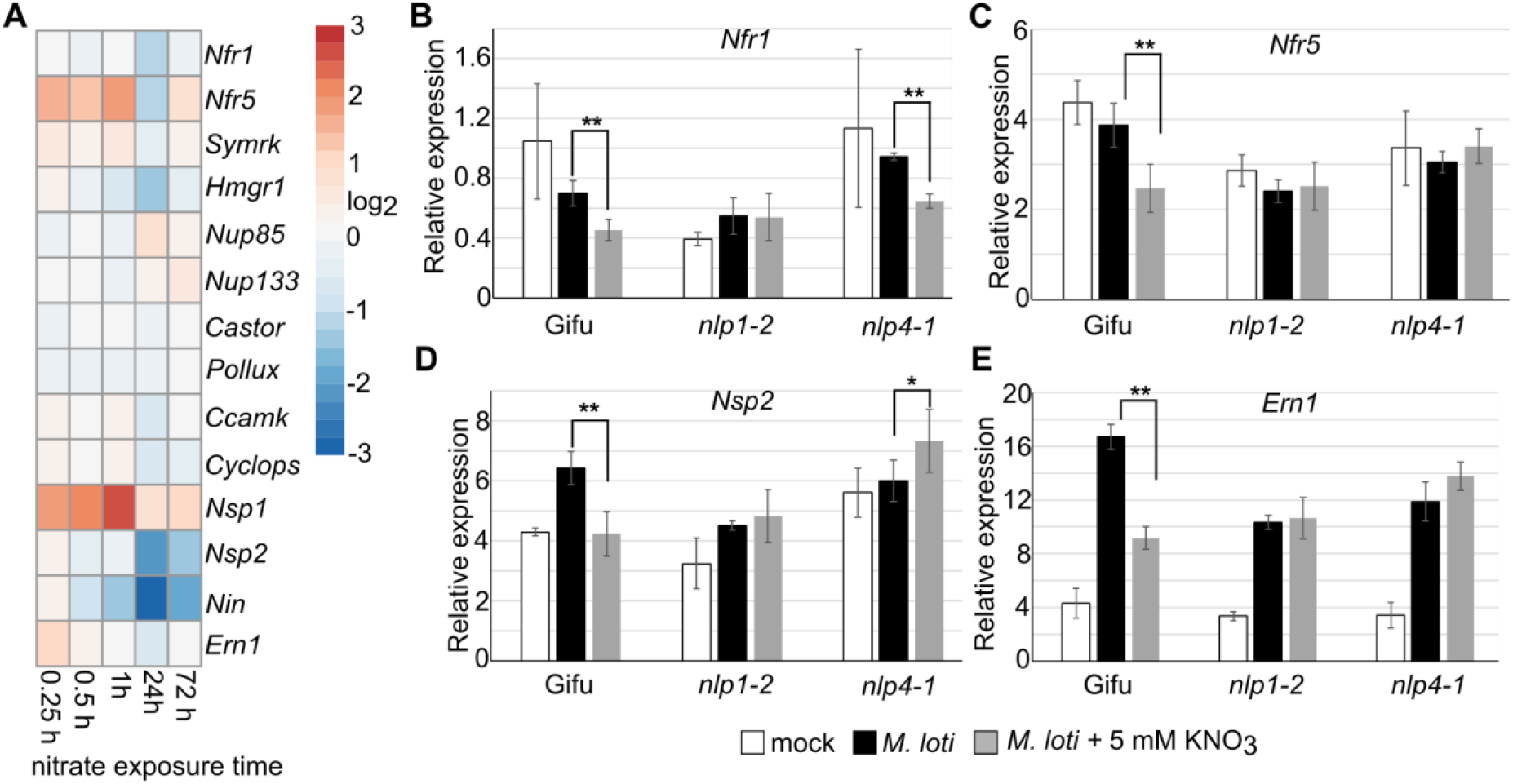
Nitrate inhibits symbiotic signalling via *Nlp1* and *Nlp4.* A, The heatmap of NF signaling gene expressions under different time series of nitrate exposure. B-E, Relative transcript abundance of *Nfr1* (B), *Nfr5* (C), *Nsp2* (D) and *Ern1 (E)* by qRT-PCR 1 dpi with *M. loti* in the absence and presence of 5 mM KNO_3_ in the indicated plant genotypes. Ubiquitin is used as a reference gene. Bars show mean +/-SE for *n*=5. Significant differences between nitrate presence and absence conditions is indicated by *<0.05 and **<0.01 as determined by Student’s t-test. Gene IDs and expression values are shown in table S1 and table S4.

## Discussion

In this study, we show that the symbiotic induction of cytokinin biosynthesis in *L. japonicus* is suppressed by high nitrate concentrations, thereby reducing the positive influence of cytokinin on nodule organogenesis. Cytokinin biosynthesis mutants, which have reduced cytokinin levels exacerbate this response and are more sensitive to nitrate inhibition of nodule development. In agreement with this, we show that this increased nitrate sensitivity can be rescued by supplementing growth media with cytokinin. We also find that *Nlp1* and *Nlp4* mediated nitrate signaling is required for this suppression of symbiotic cytokinin biosynthesis and nodule organogenesis, by suppressing the expression of upstream signalling components including *Nfr1, Nfr5, Nsp2* and *Ern1.*

### Cytokinin is a hub integrating external stimulus with nodulation signalling

In response to NF perception, many cytokinin biosynthesis genes are up-regulated in the root susceptible zone where cytokinin accumulates transiently to trigger nodule initiation (Reid et al., 2017; van Zeijl et al., 2015). The cytokinin trigger for nodulation requires signalling via the receptors (predominantly *Lhk1/Cre1)*, with loss-of-function mutants impaired in nodule development (Murray et al., 2007; Held et al., 2014; Gonzalez-Rizzo et al., 2006; Boivin et al., 2016). In contrast, cytokinin application or constitutive activation of cytokinin signaling can activate nodule organogenesis programs in the absence of rhizobia (Tirichine et al., 2007; Heckmann et al., 2011; Liu et al., 2018). Here, we find the transient induction of cytokinin in the susceptible zone is reduced when grown in high nitrate conditions. Nodule initiation and maturation, which is required for nitrogen fixation, is therefore suppressed and delayed by nitrate exposure.

In addition to the mechanism we describe where inhibited cytokinin synthesis leads to reduced nodule development, excessive cytokinin accumulation can also inhibit nodulation. This has been shown to occur through induction of Autoregulation of Nodulation (AON) and ethylene signalling, which both play negative roles in nodulation (Miri et al., 2019; Suzaki et al., 2012; Saur et al., 2011; Mortier et al., 2012; Reid et al., 2018). This is also supported by the enhanced sensitivity to nitrate inhibition of nodulation that occurs in the *Ljckx3* cytokinin over-accumulation mutants (Reid et al., 2016). We find support for this fine balance of cytokinin for nodule development with lower concentrations of cytokinin supplementation able to rescue the biosynthesis mutant impairment, or at increased levels to inhibit nodulation. Together these observations support a model where cytokinin levels are tightly regulated by multiple internal and external influences to balance the requirements for plant nitrogen between uptake from the soil and nodule development (and thus nitrogen fixation).

### Cytokinin plays a role in nitrate inhibition of organogenesis but not rhizobia infection

In contrast to the requirement in nodule organogenesis, cytokinin plays a negative role in rhizobia infection (Murray et al., 2007). Consistent with the hyperinfection phenotype of *ljlhk1*, the *ipt3-2 ipt4-1* double mutant shows more than triple the infection density of wild type. This shows that in addition to receptor signaling, cytokinin biosynthesis negatively regulates rhizobia infection. In high nitrate conditions this hyperinfection is suppressed in *ipt3-2 ipt4-1*, with a similar degree of inhibition to wild-type. This implies that at least cytokinin biosynthesis is not critical to nitrate inhibition of rhizobia infection. The ethylene and AON pathways may therefore play more prominent roles in negatively regulating infection by nitrate. In addition, *MtNlp1* may directly interfere with infection by blocking NIN’s function (Lin et al., 2018). Thus, nitrate inhibition of rhizobia infection is likely to target alternative pathways or downstream of cytokinin biosynthesis.

### *Lotus japonicus* differs from non-legumes in the cytokinin response to nitrate

In the non-legumes, cytokinin biosynthesis plays an important role in signalling root nitrate availability and coordination of shoot growth (Poitout et al., 2018; Landrein et al., 2018). This cytokinin response to nitrate has been reported in several species including *Arabidopsis* (Takei et al., 2004a), rice (Kamada-Nobusada et al., 2013) and maize (Takei et al., 2001). In response to nitrate, tZ and iP type cytokinin accumulate in shoot, but cytokinin level remains unchanged in roots of *Arabidopsis* (Poitout et al., 2018). Here we show that in *Lotus japonicus*, where cytokinin plays an important role in symbiotic organ establishment, root cytokinin biosynthesis is not enhanced by nitrate supply. Our RNA-seq data, which showed consistent induction of nitrate marker genes indicated that none of the *Ipt* or *Log* genes in *Lotus* are significantly upregulated by nitrate exposure. How *L. japonicus* signals nitrate availability relative to non-legumes may therefore differ. One possibility is that *L. japonicus* maintains a cytokinin response to nitrate in non-susceptible root tissue, with the nitrate suppressed and rhizobia enhanced cytokinin being restricted to root zones supporting nodulation. Supportive of this is that nitrate can only induce AtIpt3 in vascular bundles (Takei et al., 2004a) which differs from the predominantly cortical location of symbiotic cytokinin in *Lotus* (Reid et al., 2017). In soybean for example *GmIpt3* and *GmIpt15* are induced by nitrate (Mens et al., 2018), and the location of this induction and influence on nodule development would be interesting to determine. Both cytokinin dependent and independent pathways are thought to play roles in coordinating systemic nitrogen signals (Ruffel et al., 2011). Other root-shoot coordination pathways such as CEP signalling (Delay et al., 2013; Tabata et al., 2014) may therefore play a more prominent role in coordinating shoot growth with nitrogen availability in legumes, relegating cytokinin signalling of nitrate to a more minor role.

### *Nlp* mediated nitrate signaling inhibits NF signaling

Nitrate negatively influences many processes in nodulation, including NF signaling, nodule initiation and development, rhizobia infection and nitrogen fixation (Nishida and Suzaki, 2018). *Nlp*s play central roles in nitrate signaling and development in many species (Castaings et al., 2009; Marchive et al., 2013). Both *LjNlp4* and *MtNlp1* are involved in nitrate inhibition of nodule initiation and development, rhizobia infection and nitrogen fixation (Lin et al., 2018; Nishida et al., 2018). *LjNlp4*, in response to nitrate, directly targets *LjCle-rs2* to trigger the AON pathway and restriction of nodule initiation (Nishida et al., 2018). *MtNlp1* is able to interact and/or compete with *Nin* to block *Nin* function, which is essential for nodule initiation and development and rhizobia infection (Lin et al., 2018). We found that nitrate suppression of NF receptors expression, which is likely to reduce the susceptibility to rhizobia, requires *Nlps*. Additionally, we found *Nlp*s are also required for nitrate suppression of some nodulation transcription factors, such as *Nsp2* and *Ern1.* Taken together with previous studies, *Nlp*s regulation of nodule initiation and development, rhizobia infection and nitrogen fixation is likely to occur through restriction of key components in NF perception and signaling. This restriction results in a significant reduction in symbiotically triggered cytokinin biosynthesis which is required for nodule development. Our finding that cytokinin application can rescue the inhibitory effect of nitrate on nodule maturation and nitrogen fixation suggests that it is primarily these early signalling events that are inhibited by nitrate and not the signalling events downstream of cytokinin in nodule development. Further mechanistic studies into how *Nlp*s mediate nitrate regulation of NF signaling components and nitrogen fixation will provide opportunities to improve nitrogen fixation and yield in an economic and sustainable way.

In conclusion, we propose a model where nitrate interferes with NF signaling and symbiotic cytokinin biosynthesis in the root susceptible zone, ultimately suppressing nodule organogenesis (Fig 9). In high nitrate, *Nlp1* and *Nlp4* is activated to suppress the early symbiotic pathway components, resulting in less sensitivity to NF and reduced output of NF signaling. This reduced NF signaling capability inhibits cytokinin accumulation, which is essential for nodule initiation and development, essential prerequisites for nitrogen fixation in mature nodules.

**Figure 9.**
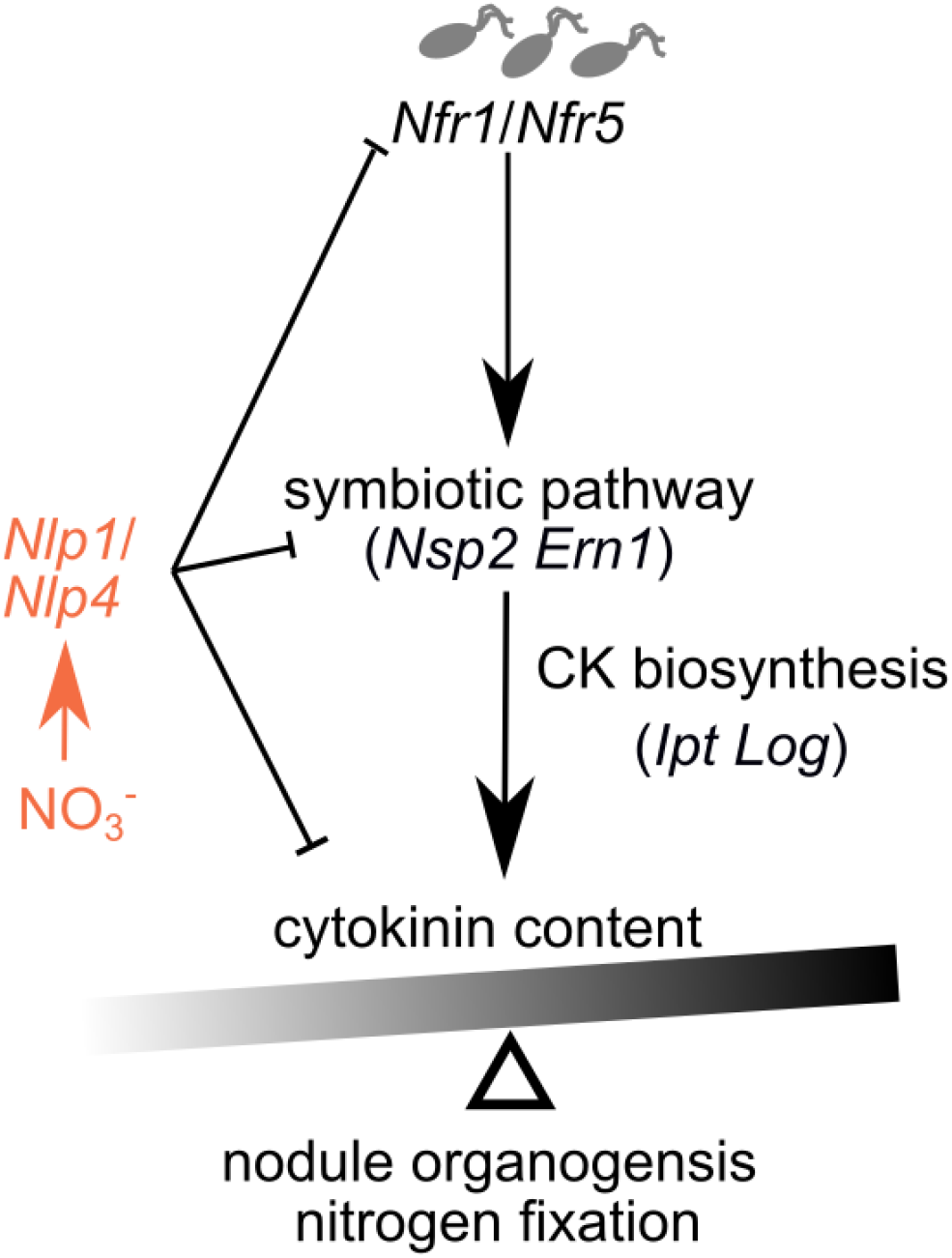
Proposed model for regulation of nodule organogenesis by nitrate. In high nitrate, nitrate signaling inhibits expression of NF receptors *(Nfr1* and *Nfr5), Nsp2* and *Ern1* mediated by *Nlp1* and *Nlp4*, therefore suppressing symbiotic cytokinin biosynthesis that is crucial for nodule organogenesis. Nodulation is thus balanced between a permissive low-nitrate state where high cytokinin levels can accumulate, or a restrictive state with low root cytokinin.

## Materials and Methods

### Plant and bacteria genotypes

*Lotus japonicus* ecotype Gifu was used as wild type (Handberg and Stougaard, 1992), while *ipt3-1, ipt4-1* and *ipt3-2 ipt4-1* were LORE1 insertion mutants (Reid et al., 2017). LORE1 mutants *nlp1-2* and *nlp4-1* were ordered through LotusBase (https://lotus.au.dk) and homozygotes were isolated for phenotyping and generation of higher order mutants as described (Małolepszy et al., 2016). Line numbers and genotyping primers are given in table S2. *Mesorhizobium loti* NZP2235 was used for nodulation assay. For infection thread observation, *M. loti* R7A strain constitutively expressing DsRed was used.

### Plant and Bacteria growth conditions

Phenotyping on growth plates was conducted by transferring 3-d old seedlings onto filter paper placed on vertical 1.4% agar noble plates containing ¼ Long Ashton (table S3) in the presence of 5 mM KCl or KNO_3_. For the cytokinin rescue assay, 6-Benzylaminopurine (BA, Sigma-Aldrich) were added into ¼ Long Ashton plates containing 5 mM KNO_3_. 3 days after transfer, seedlings were inoculated with rhizobia inoculum OD_600_=0.015. Infection threads were counted at 7 day post inoculation (dpi), while nodule numbers were counted at 14 dpi. For nitrate and RNA-seq, 3-d old seedlings were transferred on ¼ B&D plates and grown for 14 days.

### Gene expression analysis

For RNA-seq, following 14 days growth on plates, plants were acclimatised prior to treatment by submerging in ¼ B&D liquid medium overnight, then treated with 10 mM KNO_3_ for 0, 0.25, 0.5, 1, 24 or 72 hours. Root tips were removed and the remainder of the root harvested. mRNA were isolated using the NucleoSpin RNA Plant kit (Macherey-Nagel) then library preparation and PE-150 bp Illumina sequencing was conducted by Novogene.

For qPCR, the root susceptible zone was harvested at 1 or 2 dpi. mRNA were isolated using the kit described above. RevertAid Reverse Transcriptase (Thermo) was used for the first strand of cDNA synthesis. LightCycler480 SYBR Green I master (Roche Diagnostics) and LightCycler480 instrument were used for the real-time quantitative PCR. Ubiquitin-conjugating enzyme was used as a reference gene (Czechowski et al., 2005). The initial cDNA concentration of each target gene was calculated using amplicon PCR efficiency calculations using LinRegPCR (Ramakers et al., 2003). Target genes were compared with the reference genes for each of 5 biological repetitions (each consisting of 6 to 10 plants/ root susceptible zones). At least two technical repetitions were performed in each analysis. Primers used are listed in supplemental material.

### Cytokinin extraction, detection and quantification

For cytokinin extraction from *L. japonicus* material, ~20 mg of snap-frozen root and nodule material was used per sample. Samples were extracted (Floková et al., 2014) and analysed by Liquid Chromatography-Tandem Mass Spectrometry as previously described (van Zeijl et al., 2015). To determine sample concentrations, a 10-point calibration curve was constructed for each compound ranging from 1 μM to 190 pM and each dilution also contained a known amount of an appropriate deuterium-labelled internal standard.

### Acetylene reduction assay

Acetylene reduction assays were conducted essentially as described previously (Reid et al., 2016). Briefly, 250 μl air in the 5 ml glass GC vials containing the sample was replaced with 2% acetylene. Samples were incubated for 30 min before ethylene quantification using a SensorSense (Nijmegen, NL) ETD-300 ethylene detector operating in sample mode with 2.5 L/h flow rate and 6-min detection time.

### Statistical analysis

Statistical analyses were carried out using R software (R Core Team, 2015). Comparison of multiple groups included ANOVA followed by Tukey posthoc testing to determine statistical significance indicated by different letter annotations. Students t-test was used as indicated when making single comparisons.

For statistical analysis of RNAseq, A decoy-aware index was built for Gifu transcripts using default Salmon parameters and reads were quantified using the --validateMappings flag (Salmon version 0.14.1) (Patro et al., 2017). Expression was normalised across all conditions using the R-package DESeq2 version 1.20 (Love et al., 2014) after summarising gene level abundance using the R-package tximport (version 1.8.0). Differentially expressed genes with correction for multiple testing were obtained from this DESeq2 normalised data. Normalised count data for all genes is shown in supplemental table 5.

## Accession numbers

RNAseq raw data has been submitted to NCBI under BioProject accession number PRJNA642098.

## Author contributions

J.L. Conceived and performed experiments, analysed data, prepared figures and wrote the paper; Y.R. Performed cytokinin quantification; W.K. Designed cytokinin quantification procedure and analysed the data; J.S. Conceived experiments, supervised and revised the paper; D.R. Conceived experiments, analysed data, supervised and wrote the paper.

## Acknowledgements

We thank Finn Pedersen for expert plant handling and Francel Verstappen for assistance with cytokinin quantification. We are grateful for support from the project Engineering Nitrogen Symbiosis for Africa (ENSA) currently supported through a grant to the University of Cambridge by the Bill & Melinda Gates Foundation and UK government’s Department for International Development (DFID). Analysis by Y.R. and W.K. is supported by Netherlands Organization for Scientific Research (VENI863.15.010).

